# Intersecting effects of social circumstances and transcendent thinking on mid-adolescents’ longitudinal functional connectome development

**DOI:** 10.1101/2025.05.20.654735

**Authors:** Amir Hossein Ghaderi, Xiao-Fei Yang, Rebecca Gotlieb, David L. Johnson, Mary Helen Immordino-Yang

## Abstract

Socially moderated variation in mid-adolescents’ functional brain network (FBN) development is insufficiently studied, especially within low-SES contexts. Alongside demographic factors, evidence suggests adolescents’ malleable psychological dispositions may contribute, with possible clinical and educational implications. We applied graph theory to a unique 2-year longitudinal fMRI dataset (N=65) with an ecologically valid 2-hour interview revealing an emergent mid-adolescent psychological disposition, “transcendent thinking” (TT)—adolescents’ tendency to consider systems-level, ethical, personal implications of social information. Overall, FBNs showed increased segregation and entropy but decreased integration and energy. Modularity increased particularly in somatosensory subnetworks. Classifying participants by two key demographic factors, parents’ education (PE) and community violence exposure (CVE), FBN changes differed by group, and TT moderated changes in high-CVE/low-PE participants, arguably the most vulnerable group. Machine learning revealed TT and CVE, but not IQ, as principal FBN change predictors across groups. Findings suggest social context and psychological dispositions influence low-SES adolescents’ FBN development.

## 1– Introduction

Mid-adolescence, here defined as age 14-18, is a critical phase of social sensitivity in human brain development^1^. During this period, developmental transformations occur in inter-regional functional connectivity, resulting in the refinement of functional brain networks (FBNs)^1–3^. These transformations vary significantly across individuals^1,2^. Variation may in part relate to social circumstances, such as exposure to violence^4–7^ and various challenges associated with low socioeconomic status (SES)^8–11^. In particular, adolescents living in low-SES urban communities are exposed to varying degrees to social challenges that may have implications for their FBN development. At the same time, developmental research suggests resilience in the form of positive psychological factors, such as malleable “ways of thinking”, that may also influence brain development^7,12,13^, even beyond the effects of IQ^14^, and may possibly mitigate the extent to which negative demographic circumstances affect the development of FBNs. This issue has important implications for our understanding of adolescent brain development in low-SES contexts, and for clinical and educational practice designed to support youths. Longitudinal studies are much needed, especially studies that leverage methodological advances for analyzing FBNs in mid-adolescent populations in low-SES contexts and include rich psychological information about participants.

Given the above, here we studied the intersecting effects of key demographic and psychological factors on developmental variability in longitudinal change in mid-adolescent FBNs within a low-SES community sample. 65 healthy, non-delinquent mid-adolescents from stable family situations, aged 14-18 years at the start of the study, underwent two resting state functional neuroimaging sessions roughly two years apart. Our dataset includes two demographic factors highlighted in previous studies^12,15^ as influencing neural development: parental education (PE) levels and community violence exposure (CVE), as positive and negative influences respectively. It also includes an ecologically valid measure of a malleable psychological capacity, transcendent thinking (TT), that serves as a superordinate construct that integrates findings from a century of developmental psychological science related to healthy psychosocial development in mid-to lateadolescence^16–19^. In TT, adolescents leverage their emerging capacities to consider complex systems-level, ethical and personal implications of the information they learn, especially within social contexts, “transcending beyond” the concrete, specific and context-dependent aspects. Engaging in TT has been shown to moderate the activation and deactivation of key large-scale FBNs, among them the salience (SN), default-mode (DMN) and frontoparietal (FPN) networks^12,15^, and to show longitudinal changes in functional connectivity among specific networks, with psychosocial implications in young adulthood^12^. Including this ecologically valid measure in a graph theoretical analysis (GTA) allows us to investigate the possible roles and interactive effects of important demographic and psychological factors on the development of FBNs in mid-adolescents.

Previous FBN studies have been predominantly cross-sectional and have not specifically addressed the mid-adolescent age range or involved youth from low-SES communities. These studies have shown group-level developmental changes in FBNs^20–23^ marked by increased modularity and segregation with age; such results have been replicated across various analytical and methodological approaches^12,24–32^. Additionally, many of these studies have reported increased global connectivity at the whole-FBN level or between specific modules, often interpreted as enhanced integration^33^. Studies of FBN development at the level of modules have also indicated substantial changes in both intra-network and inter-network functional connectivity. These changes are predominantly characterized by increased modularity in networks such as the DMN^26,34^, SN^35^, FPN^2^, and sensory networks^30^. A recent study with a wide age range from childhood to late adolescence highlighted significant developmental changes in the sensorimotor network (SMN), and proposed that this may serve as a foundational basis for other modular changes^36^.

While these findings are promising, the rapid advancements in network analysis and novel approaches in GTA methods underscore the necessity for further, more in-depth investigations into the effects shown in these previous studies, and in particular to examine sources of variation and resilience within low-SES groups. These advancements are needed to gain a better understanding of FBN in adolescents from two perspectives. First, especially in the context of ecologically valid psychological information, the application of new GTA measures opens novel avenues for research. Notably, in recent years, findings from spectral GTA (sGTA) have shown that, beyond network topology, the energy of FBNs (related to stability of synchrony) can be influenced by conditions related to aspects of health and disease^37–41^. Studies have also shown that information entropy (Shannon entropy) can index complexity of network connectivity^38,40^. Examining the energy and entropy trajectories of FBNs in adolescents, and relating these to social circumstances and psychological dispositions, could provide insights into the underlying mechanisms shaping FBN development.

Second, the omission of null models in previous investigations makes it challenging to determine whether the observed increases in network properties such as integration and segregation result from a general increase in functional connectivity across the entire network or from specific rearrangements during development in functional circuits responsible for neural information propagation^42^. This issue also applies to the characteristics of modules. In many previous studies, it is unclear whether the increase in connectivity within a module, typically interpreted as an increase in the modularity of that module, is due to a general increase in the connectivity weights across the entire network, or to an increase just within that module. Here, we first applied null models to examine the overall network characteristics (including topological and spectral measures) and then, using data-driven methods^43–45^ and innovative approaches for applying null models, investigated the modularity ratio and the relationships of modules with other components. This combination of analyses, along with the nature of the dataset, provides a unique and generative contribution.

To conduct our study, we initially analyzed the network-wide and modular features at the whole-sample level to examine longitudinal changes shared across participants, to characterize topology, dynamics, and information processing in the FBNs, and to investigate whether the findings would differ with the application of null models. We then investigated the effects of demographic factors on the development of FBNs. Building on existing evidence of experience-related effects on brain development, we applied an unsupervised clustering method based on the two demographic factors, CVE and PE. This approach enabled us to categorize our participants into three distinct groups: 1) high-PE/low-CVE, 2) low-PE/low-CVE, and 3) low-PE/ high-CVE, and to characterize and compare these groups’ FBN development.

Next, using the data grouped by demographic factors, we examined how psychological factors influenced the development of global and modular FBN features in each group, given recent evidence that TT is a potential marker of longitudinal change in functional/structural brain development^7,12^. To confirm and extend our analyses, we then used a supervised support vector machine (SVM) model to predict various FBN features at the second brain scan session for each participant, using as inputs combinations of FBN features from the first scan session (conducted two years earlier), as well as demographic and psychological variables and age. We determined the relative contributions of each input parameter to predicting the network features in the second session. We hypothesized that not simply demographic but also psychological factors would be predictors of FBN longitudinal change, and that the models’ accuracy would be lower in the Low-PE group due to the increased complexity/risk associated with these participants’ social circumstances.

*Study Importance*: Our study is important for several reasons. We utilized longitudinal data from a modestly scaled project designed to capture a cohort of mid-adolescents’ psychosocial development with unprecedented ecological validity and psychological nuance^12,15^. There is an urgent need in the psychological and brain sciences to study populations that have not traditionally been involved in research, including non-White individuals living in low-SES communities, and in particular to focus on factors that support these populations’ normative development, rather than solely focusing on deficits and risks without also considering endogenous factors that promote resilience. Related, there is a dearth of studies examining the effects of malleable, psychosocial developmental factors, such as habits of mind, that can be strategically supported by developmental interventions in education or outside of school^13^. Our sample reflects the demographic diversity present in many urban low-SES communities; participants’ parents had immigrated to the United States from thirteen different countries, primarily in East Asia and Latin America. Asian and Latinx youth are substantial and growing segments of the U.S. population. Our sample is also reflective of the variation in parental education levels and financial circumstances that exists within low-SES communities.

## 2– Results

Fig. 1 shows the analytical pipeline to construct FBNs and evaluate their properties. We performed whole-group level analyses as well as analyses of separated groups based on the demographic and psychological measures. In the first phase of the study, we performed group analyses and investigated longitudinal changes of network measures between neuroimaging sessions.

**Fig. 1:**
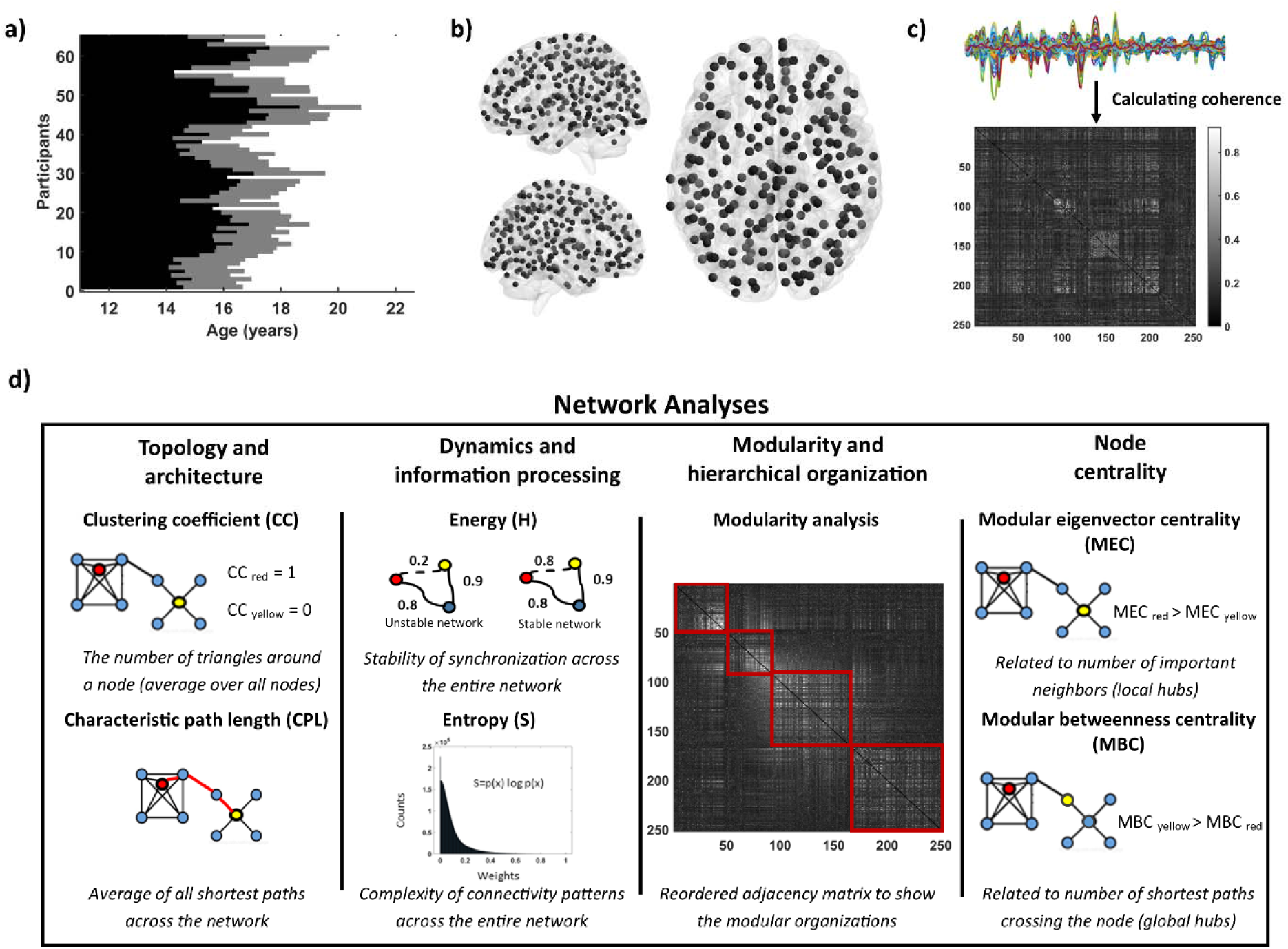
Methodological pipeline. a) A longitudinal resting state fMRI dataset involving a cohort of 65 individuals (N=65, f=36) during their adolescent years (aged 14-18 years at recruitment) was used. This dataset was collected across two recording sessions, with a subset of participants failing to complete fMRI recording in the second session. Ages and recording sessions for each participant are visualized. b) Upon preprocessing the data, we used a well-established anatomical atlas featuring 264 distinct brain regions of interest (ROI)^46^. Given the presence of inadequate BOLD signal quality within 12 regions across various participants, we excluded these regions. c) Coherence between all possible pairs of 252 ROIs was calculated. These coherence metrics were computed for each participant within each session and then used to construct adjacency matrices (by considering each brain region as a node and the coherence between each pair of nodes as an edge) which represent functional brain networks. d) To probe the topology, dynamics, and complexity of functional connectomes, we harnessed various approaches in conventional and spectral graph theoretical analysis, and information theory. Our assessment of these networks unfolded across three tiers: firstly, the entire brain networks (four measures of CC, L, H, and S); secondly, modular functional structures; and lastly, regional connectivity patterns.

### 2-1 Group level analysis

#### 2-1-1 Whole-FBN features

As shown in Fig. 2a, the connection density (CD; defined as average functional connectivity over whole-FBN) is stronger in Session 2 (S2) compared to session 1 (S1). Statistically, the permutation t-test with false discovery rate (FDR) correction revealed a significant difference between S1 and S2with lower values in S1 in CD (t=-3.81, q=0.0046). Before applying null models, lower clustering coefficient (CC) (t=-3.68, q=0.006), energy (H) (t=-6.34, q<0.001), and entropy (S) (t=-4.37, q=0.001), as well as higher characteristic path length (CPL) (t=3.85, q=0.005), were observed in the S1 (Fig. 2b).

**Fig. 2:**
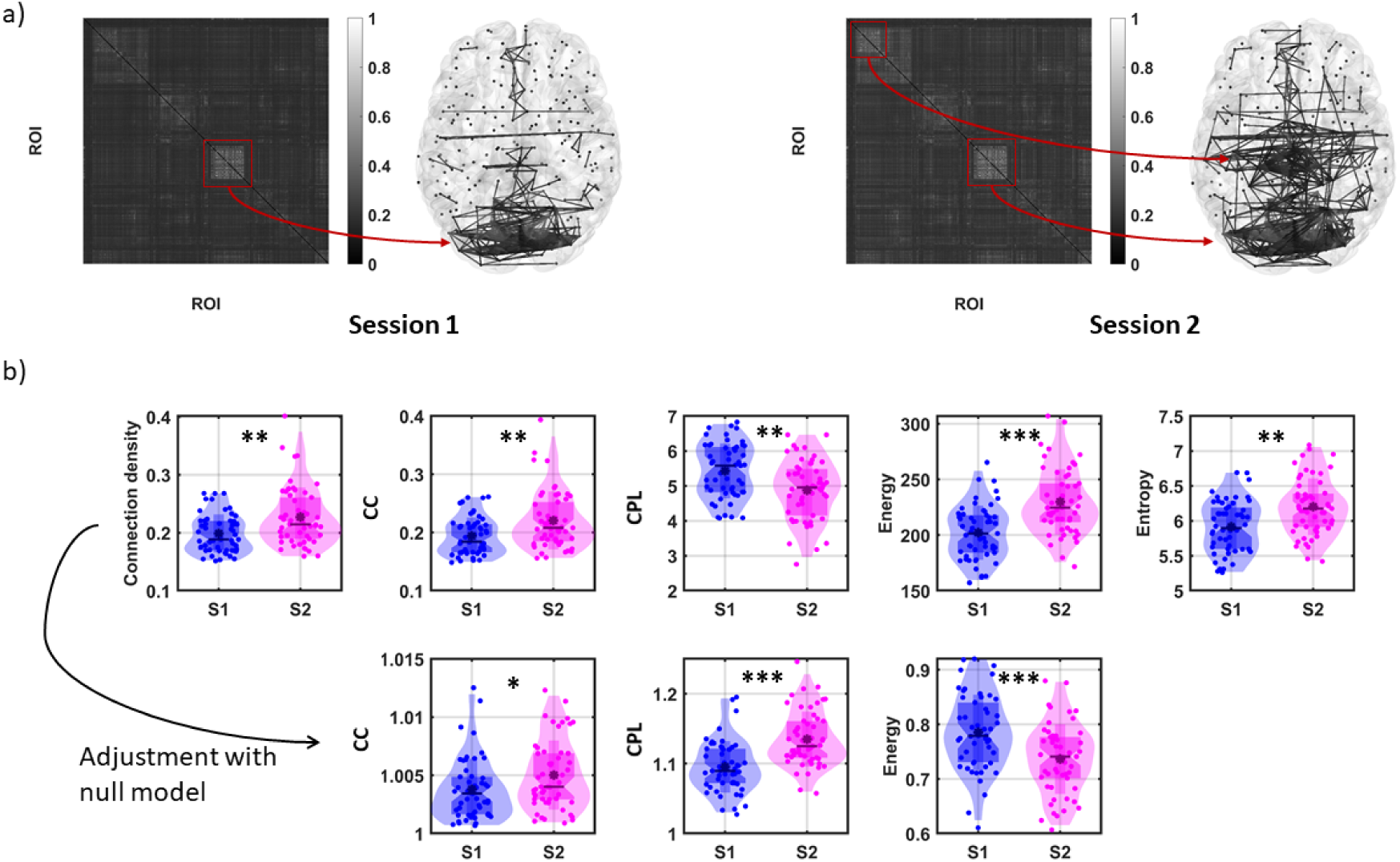
Functional connectivity maps and whole-FBN features: a) Adjacency matrices of weighted networks and the strongest connectivity patterns in both sessions. In these matrices, each row and column represent a specific brain region, and the color at the intersection of a row and column indicates the strength of connectivity between the corresponding brain regions. To highlight the strongest connectivity patterns in each session, we applied a sparsity approach and set a threshold to visualize the most robust connections. In Session 1, the strongest connections are observed within the visual network. In Session 2, the strongest connections are observed in both the visual and sensorimotor networks. b) Comparison of five whole-FBN measures between the two sessions. Raw values of whole-FBN measures are presented at the top row and the corrected (using null models) whole-FBN measures are shown at bottom. Importantly, the developmental direction of characteristic path length and energy are changed after applying null model corrections. * p < 0.05, ** p < 0.01, *** p < 0.005.

Next, CC, CPL, and H were calculated against randomly shuffled null models (CD and S are not sensitive to the location of connections in the network and remain equal in the random-shuffled null and original models). The CC results were consistent after applying null models, with a significantly lower value observed in the S1 compared to the S2 (t=-2.52, q=0.015). However, the CPL and H showed opposite results after applying null models, with a significant lower value in CPL (t=-6.10, q<0.001) and a higher value in H (t=4.41, q<0.001) in the S1 compared to the S2 (Fig. 2b).

Given current calls in GTA to apply null models^42^, we used the values normalized against null models for all remaining analyses in the study. To perform longitudinal analyses and retain participants with only one neuroimaging session, we utilized linear mixed models. This approach enabled us to account for variations in age and in the intervals between successive scans. Our model included FBN features as the outcome variable, with age (linear term) as a fixed factor and participants as a random factor (i.e., FBN feature = age + (1|participant)). FDR analyses showed significant longitudinal change of FBN on all five features: CD (t=3.62, q=0.004), CC (t=3.48, q=0.007), CPL (t=5.82, q<0.001), H (t=-4.61, q<0.001), and S (t=4.22, q<0.001). The results were consistent with results of the permutation t-test between two sessions.

#### 2-1-2 Modularity analysis

As shown in Fig. 3-a, modularity analysis demonstrated that the optimal modularity (with less than 10% isolated nodes) of FNBs occurs at a modularity parameter γ=1.03. At this γ, the FBNs are divided into five distinct modules (Fig. 3-b). The degree of overlap between these modules and predefined FBNs^46^ is illustrated in Fig. 3-c; a comprehensive description of these subnetworks is presented in Fig. 4.

**Fig. 3:**
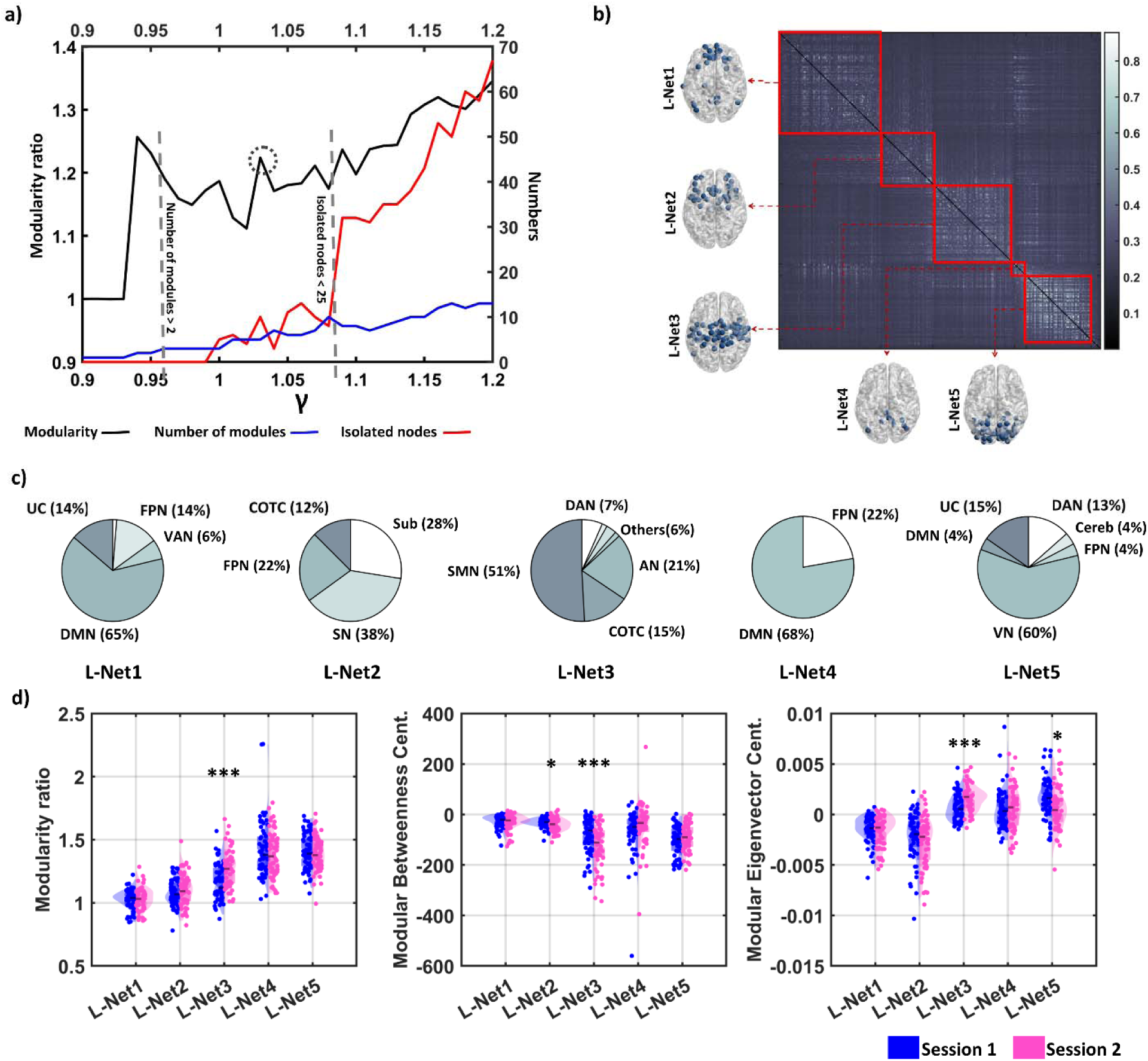
Modularity analysis (large modules). a) A hierarchical algorithm^43,44^ was applied to systematically evaluate the modular configuration inherent within the FBNs. The outcome of this partitioning process depends on the modularity parameter, denoted as γ. An increase in γ leads to the subdivision into smaller subnetworks (resulting in more subnetworks). To determine the optimal γ value, we calculated the modularity ratio of each module. Then, the modularity ratios of subnetworks were averaged (a higher average modularity value indicates the optimal γ value). As γ increases, the number of isolated nodes also increases. To avoid excessive isolated nodes, we set a threshold at 10% of all nodes (i.e., 25). The maximum modularity ratio while the number of isolated nodes remained below 10% was achieved at γ = 1.03. b) The modularity analysis identified five distinct modules (L-Net1 to 5). c) The configuration of each module was compared with the modules in the Power et al. (2010) study. Each module is presented with the percentage of nodes assigned to different modules in the Power et al. (2010) study. DMN: default mode network, FPN: frontoparietal network, VAN: ventral attention network, UC: uncertain, SN: salience network, Sub: subcortical, COTC: Cingulo-opercular task control, SMN: Sensory/somatomotor network, AN: auditory network, VN: visual network, DAN: dorsal attention network, and Cereb: cerebellum. d) Left: Modularity ratio measures the average connectivity between nodes in one module divided by average connectivity in a randomly shuffled network with same nodes. A significant difference between two sessions was observed in L-Net3 Middle: Modular betweenness centrality measures the average shortest paths that cross the regions in a module, normalized against a randomly shuffled network. Significant differences between sessions were observed in L-Net2, and L-Net3. Right: Modular eigenvector centrality measures the average local connectivity of regions within the module (compared to the null model). Significant differences between sessions were observed in the L-Net3, and L-Net5. *p < 0.05, **p < 0.01, ***p < 0.005.

**Fig. 4:**
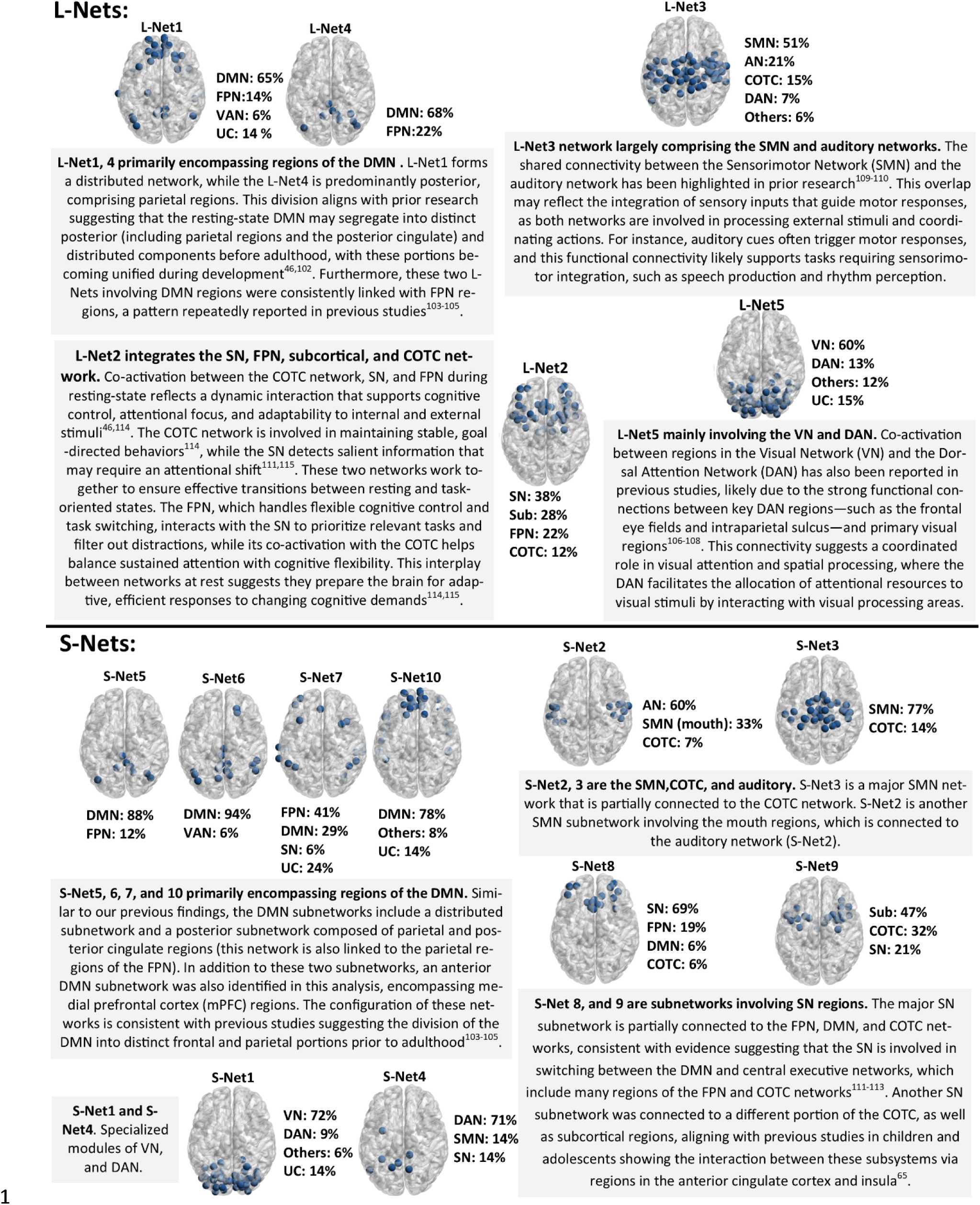
Configuration of modules. The architecture and configuration of both L-Net and S-Net sets are explained and interpretated. Separation and merging in subnetworks were explained and discussed according to previous studies^46,65,102–115^.

Then, as shown in Fig. 3-d, we calculated modularity ratio (MR), modular betweenness centrality (MBC), and modular eigenvector centrality (MEC). In comparing the S1 and S2 statistically, permutation t-tests and FDR revealed significant lower value in MR in S1 in L-Net3 (t=-4.76, q<0.001), which primarily consists of the SMN and auditory network. The analysis of MBC showed a general trend toward negative values for most participants, indicating that the shortest paths between regions in the FBN are longer than those in the randomized null models. Permutation t-tests and FDR revealed significant higher value in MBC in the S1 in L-Net2 (t=2.47, q=0.042), which primarily consists of the SN, subcortical, and FPN regions, and again in L-Net3 (t=4.20, q<0.001). Finally, MEC showed significant lower value in L-Net3 (t=-4.33, q<0.001) and higher value in L-Net5 (t=2.48, q=0.041) (primarily consists of the visual network) in S1.

To segregate modules into more specific groups of regions, we also analyzed higher γ values using a more flexible criterion that allowed for 20% isolated nodes. With this criterion, the maximum MR was achieved at γ=1.15 (Fig. 5-a), resulting in the identification of 10 modules (S-Net1 to S-Net10) (Fig. 5-b). The contribution of these modules to the involvement of FBNs is illustrated in Fig. 5-c. The configuration of these networks is explained in the Fig. 4.

**Fig. 5:**
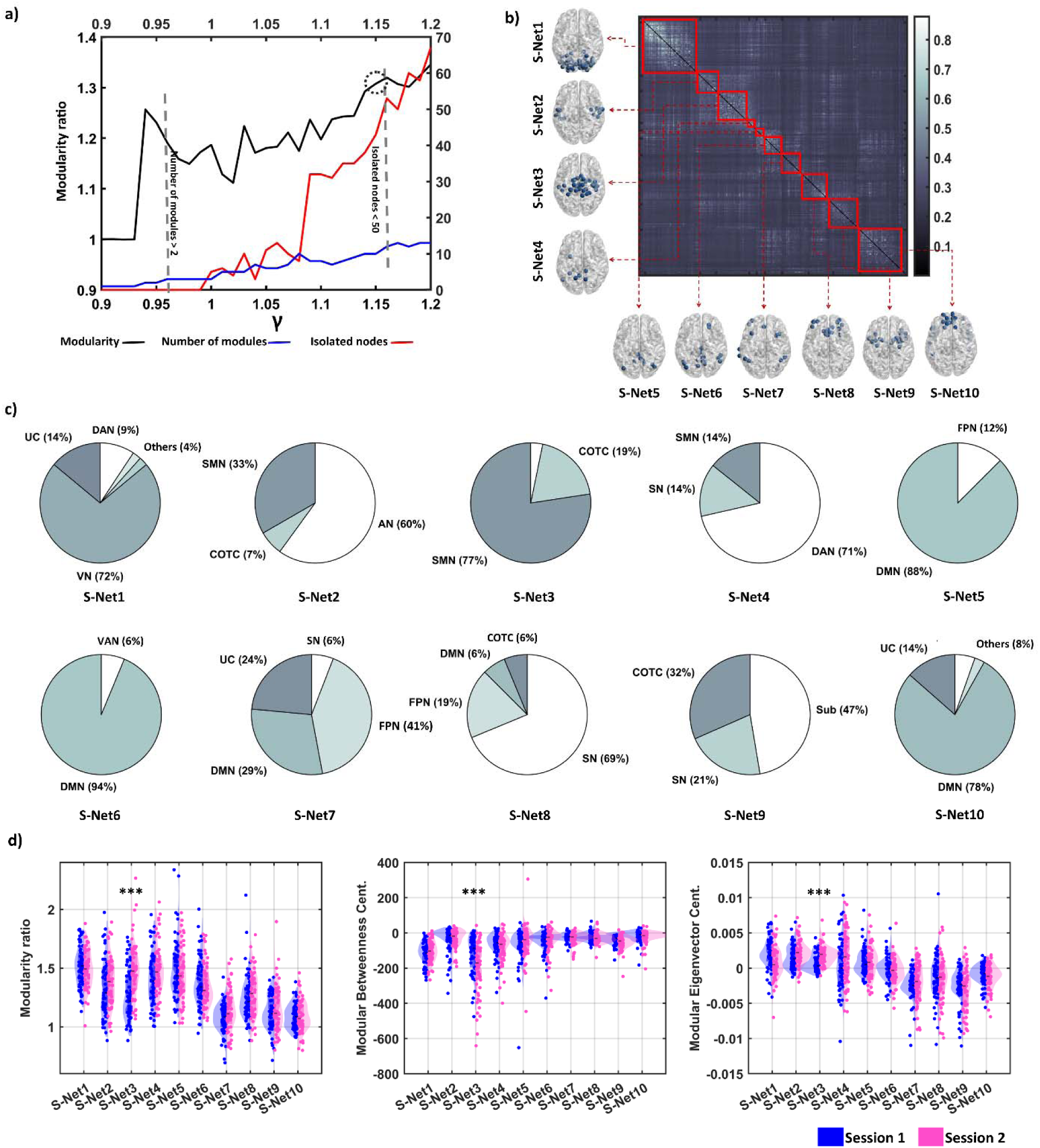
Modularity analysis (small modules). a) A hierarchical algorithm^43,44^ was applied to systematically evaluate the modular configuration inherent within the FBNs (see the caption of Fig. 3 and the method section for more details). To achieve smaller modules, we set a threshold at 20% of all nodes (i.e., 50) as the acceptable number of isolated nodes. The maximum modularity ratio while the number of isolated nodes remained below 20% was achieved at γ = 1.15. b) The modularity analysis identified ten distinct modules (S-Net1 to 10). c) Again, each module is presented with the percentage of nodes assigned to different modules in the Power et al. (2010) study. DMN: default mode network, FPN: frontoparietal network, VAN: ventral attention network, UC: uncertain, SN: salience network, Sub: subcortical, COTC: Cingulo-opercular task control, SMN: Sensory/somatomotor network, AN: auditory network, VN: visual network, DAN: dorsal attention network, and Cereb: cerebellum. d) **Left**: Modularity ratio measures the average connectivity between nodes in one module divided by average connectivity in a randomly shuffled network with the same nodes. A significant difference between two sessions was observed in S-Net3 **Middle**: Modular betweenness centrality measures the average shortest paths that cross the regions in a module, normalized against a randomly shuffled network. A significant difference between sessions was observed in S-Net3. **Right**: Modular eigenvector centrality measures the average local connectivity of regions within the module (compared to the null model). A significant difference between sessions was observed in S-Net3. *p < 0.05, **p < 0.01, ***p < 0.005.

Permutation t-tests and FDR comparing both sessions revealed significant lower value in S-Net3 (primarily consisting of the SMN) in MR (t=-5.76, q<0.001), and MEC (t=-5.36, q<0.001), but higher value in MBC (t=4.41, q<0.001) in S1. These results align with our previous results involving five larger networks (L-Nets) (Fig. 5-d).

### 2-2 Groups based on PE and CVE

#### 2-2-1 Whole-FBN features

To investigate the effects of two key demographic factors, i.e., CVE and PE, on FBN development across individuals, we first conducted k-means clustering analysis, optimized using the Calinski–Harabasz index, to determine how participants could be grouped based on these variables. The results indicated that the optimal number of clusters was three, which comprised: (1) a group with high-PE/low-CVE, (2) a group with low-PE/low-CVE, and (3) a group with low-PE/high-CVE (Fig. 6-a).

**Fig. 6:**
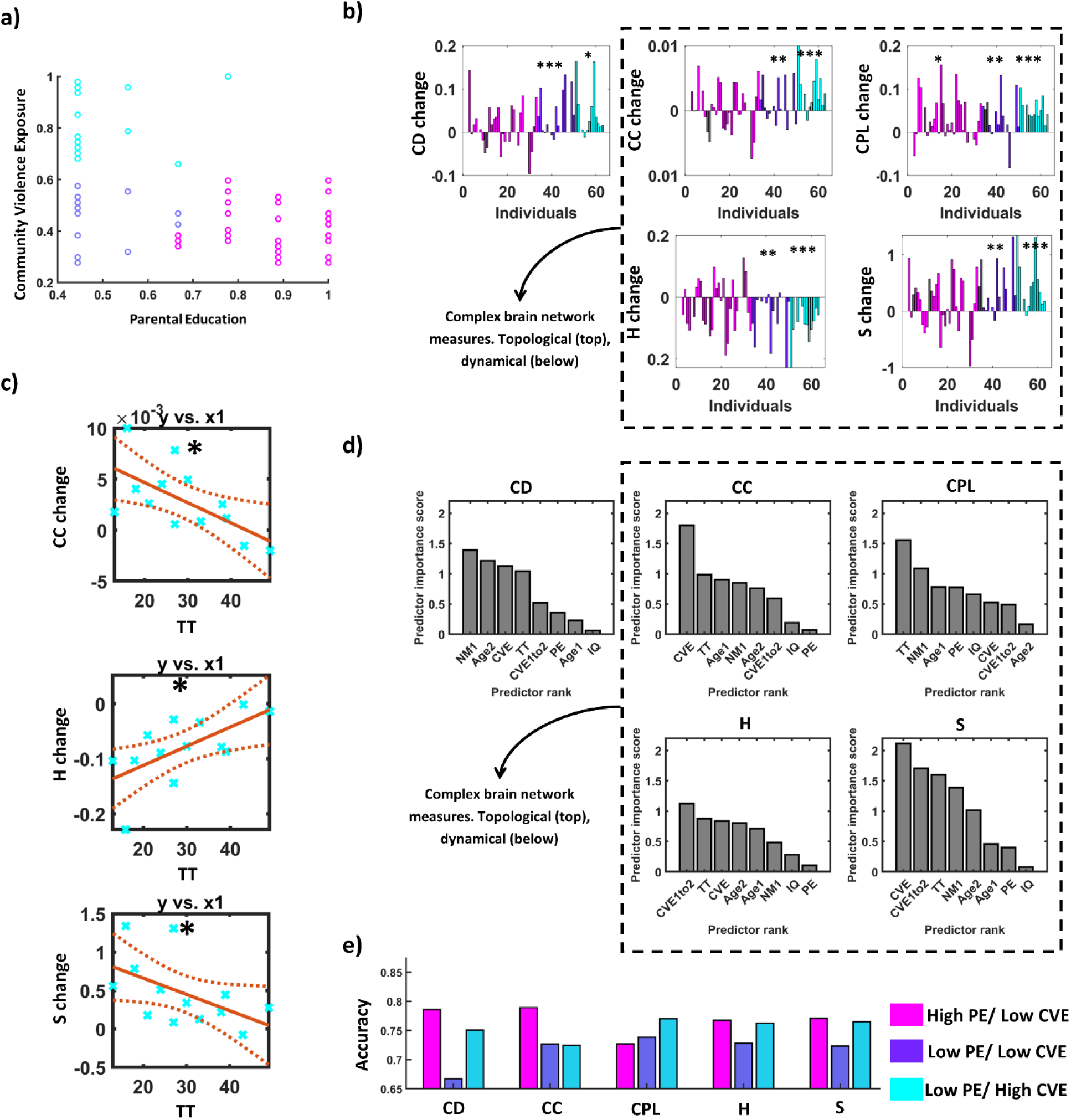
Classification, group level, and machine learning analysis. a) K-means analysis divided our samples into three groups: 1) high PE/low CVE, 2) low PE/low CVE, and 3) low PE/high CVE. b) Significant longitudinal changes in FBN measures (for both connection density and complex network measures) were observed for the two groups with low PE. The group with high PE showed a significant longitudinal change only in characteristic path length. c) Partial correlation analyses (controlling for age) showed that participants with higher transcendent thinking scores exhibited less change in the three network measures. This effect was observed only in the group with low PE/high CVE. *p < 0.05, **p < 0.005, ***p < 0.001. d) A machine learning approach was used to predict the FBN features from a set of input parameters. F-test analysis ranked the predictors for each FBN measure. For the complex network measures, CVE and TT were consistently the most important predictors. e) The accuracy of a support vector machine (SVM) model is presented for various FBN measures and groups. Overall, the SVM model predicts more accurately in the group with high PE/low CVE. TT: transcendent thinking, CT: concrete thinking, PE: parental education, CVE: childhood and early adolescent community violence exposure, CVE1to2: mid-adolescent community violence exposure CD: connection density, CC: clustering coefficient, CPL: characteristic path length, H: energy, S: entropy.

We then performed a longitudinal analysis using linear mixed models to assess developmental changes in FBN measures within each group. The results revealed that the group with high-PE/low-CVE exhibited a significant change in topological brain integration, as indicated by an increase in CPL over time (t=3.113, q=0.003), suggesting decreased brain integration with age in this group (Fig. 6-b). In contrast, the other two groups showed significant changes in additional whole-FBN features. The group with low-PE/low-CVE displayed significant changes in density (t=3.670, q=0.001) CC (t=2.791, q=0.010), CPL (t=3.225, q=0.004), H (t=-3.926, q<0.001), and S (t=4.131, q<0.001). Similarly, the group with low-PE/high-CVE exhibited significant changes in CC (t=3.128, q=0.004), CPL (t=7.530, q<0.001), H (t=-4.501, q<0.001), and S (t=3.042, q=0.005) (Fig. 6-b).

Finally, we conducted partial correlation analyses (age was controlled) between changes in FBNs and each psychological component. Following FDR correction, significant correlations were observed between TT and changes in three network features; CC (R=-0.65, q=0.022), H (R=0.69, q=0.014), and S (R=-0.59, q=0.043), only in the group with low-PE/high-CVE (Fig. 6-c).

#### 2-2-2 Machine learning analysis

To further validate our previous findings and assess the effects of various demographic and psychological measures on predicting developmental patterns for each individual, we applied a *F*-test feature selection analysis followed by a supervised SVM analysis. This additional analysis was used to predict whole-FBN measures (i.e., CD, CC, CPL, H, and S) in the second session by using FBN measures in the first session, age at both sessions, psychological/demographic (TT, IQ, PE, CVEs) metrics as inputs. Overall, as shown in the Fig. 6-d, the *F*-test results showed that the TT is an important predictor for predicting all four complex brain network features (i.e., CC, CPL, H, and S). CVE was also an important predictor for complex brain network features except for CPL. The SVM analysis revealed a good accuracy of prediction for FBN features in the second session (Fig. 6-e). The highest prediction accuracy was achieved for the group with high-PE/low-CVE, suggesting a smoother and more predictable developmental trend for participants in this group.

#### 2-2-3 Modularity analysis

We then applied linear mixed models to analyze the longitudinal changes in MR, MBC, and MEC across three groups. In general, we observed changes in the three modules. 1) SMN and auditory network, 2) SN, subcortical, and COTC, and 3) DMN.

In the SMN and auditory module, all three groups exhibited longitudinal changes in both L-Net and S-Net analyses. However, unlike the other two groups, the low-PE/high-CVE group did not show any significant changes in MEC over L-Net3 and S-Net2. Additionally, only high-PE/low-CVE group showed significant change in MBC over S-Net3.

In the SN, subcortical, and COTC module, the low-PE/high-CVE group exhibited a decrease in MEC over both small and large modules. The low-PE/low-CVE group showed this decrease only in the small module, while the high-PE/low-CVE group did not display any changes in this module.

Finally, regarding the DMN module, the high-PE/low-CVE group was the only one that demonstrated a decrease in MEC over a small module. The statistical details of the significant results are provided in Table 1.

**Table 1:** Modularity changes across groups. Linear mixed models and FDR revealed distinct significant developmental changes of L-Net and S-Net modules across groups.

| SMN and auditory network | MR | MBC | MEC |
| --- | --- | --- | --- |
| **L-Net3** |  |  |  |
| High PE/Low CVE | *↑ t=2.868, q=0.028 | *↓ t=-3.056, q=0.017 | *↑ t=2.836, q=0.031 |
| Low PE/Low CVE | *↑ t=4.454, q<0.001 | *↓ t=-3.050, q=0.028 | *↑ t=4.363, q=0.001 |
| Low PE/High CVE | *↑ t=2.927, q=0.034 | *↓ t=-2.770, q=0.049 | - |
| **S-Net2** |  |  |  |
| High PE/Low CVE | - | - | *↑ t=2.849, q=0.048 |
| Low PE/Low CVE | - | - | *↑ t=2.980, q=0.042 |
| Low PE/High CVE | - | - | - |
| **S-Net3** |  |  |  |
| High PE/Low CVE | *↑ t=3.592, q=0.013 | *↓ t=-3.231, q=0.020 | *↑ t=2.965, q=0.044 |
| Low PE/Low CVE | *↑ t=3.908, q=0.006 | - | *↑ t=3.396, q=0.022 |
| Low PE/High CVE | *↑ t=2.957, q=0.045 | - | *↑ t=3.176, q=0.037 |
| SN, Subcortical, and COTC | MR | MBC | MEC |
| **L-Net2** |  |  |  |
| High PE/Low CVE |  |  |  |
| Low PE/Low CVE |  |  |  |
| Low PE/High CVE |  |  | *↓ t=-2.941, q=0.038 |
| **S-Net9** |  |  |  |
| High PE/Low CVE | - | - |  |
| Low PE/Low CVE | - | - | *↓ t=-2.954, q=0.045 |
| Low PE/High CVE | - | - | *↓ t=-2.942, q=0.049 |
| DMN | MR | MBC | MEC |
| **S-Net6** |  |  |  |
| High PE/Low CVE |  |  | *↓ t=-2.806, q=0.041 |
| Low PE/Low CVE |  |  |  |
| Low PE/High CVE |  |  |  |

Our partial correlation analysis, adjusted with FDR multiple comparison correction, revealed no significant relationships between demographic and psychological measures and changes in MR, MBC, and MEC (session2 minus session1) in all groups.

## 3– Discussion

This study employs comprehensive analyses from graph theory, information theory, and statistical and machine learning models to explore the longitudinal development of FBNs in a mid-adolescent, diverse low-SES sample whose development of a key mid-adolescent psychosocial milestone, TT, was characterized out the outset.

Across the sample, after applying null models, we observed significant changes in FBNs over the two-year period, including decreased brain integration (inversely related to characteristic path length) and increased modularity across major networks, most consistently in the SMN. Notably, change over time in other FBN features (e.g., S, H and CC) varied with respect to demographic and psychological factors. TT and CVE were important predictors of developmental FBN change, while IQ was not. TT was an especially important moderator of FBN change among youth with high CVE.

### 3-1 All-sample analysis

#### 3-1-1 Whole-FBN measures

Our initial analysis using GTA showed that both segregation and integration in FBNs, at the whole-brain scale, increase with age during mid-adolescence. These findings were based on calculating the CC (a measure of segregation) and CPL (a measure of integration). These results are consistent with previous GTA and component analysis studies, which suggest an increase in both FBN integration and segregation with age during adolescence^22,30,33,46^. However, the increase in integration contradicts some earlier findings, which suggested that interconnectivity between subnetworks decreases with age, implying a reduction in integration^22,30,46^. Consistently, after applying null models, results revealed that although the brain segregation increases with age, a reduction in FBN integration is observed. This result can suggest that our initial observation of increased FBN integration (before applying null models) was due to an overall increase in connectivity strength with age, rather than to a topological reorganization of the FBN to create shorter pathways for neural information processing across the entire network. Thus, our analysis demonstrates that while the brain generally strengthens connectivity between different regions with age during adolescence, this increase appears not to result in shorter pathways between distant regions. Instead, the brain tends to increase the length of pathways between functional far regions. These findings align with earlier studies that analyzed connectivity pathways between and within modules without applying graph theory measures^22,46^.

Our findings from sGTA and information theory analyses (i.e., calculating H and S) provide additional insights, but these results were achieved for the first time in this study and cannot be directly compared to previous studies. Our initial H results showed an increase in H, which could suggest a longitudinal change in network synchronization stability^38^. However, similar to brain integration, this result could simply be a consequence of an increase in the average connectivity weights across the entire network, rather than reflecting changes in the stability of synchronization or the dynamics of connections between regions. After applying null models, the results showed a reduction in H, confirming that the initial finding was due to increasing average connectivity weights. Conversely, the stability of synchronization is reduced across the entire network, which aligns with our entropy results showing an increase in entropy with age, consistent with the notion that the brain increases diversity and complexity of connectivity across mid-adolescence.

#### 3-1-2 Modularity measures

A description of the modular functional architecture of L-Nets and S-Nets is presented in Fig. 4. Among all networks, L-Net3 (encompassing the SMN and auditory networks) showed a highly significant increase in the MR and MEC, combined with a significant decrease in MBC. This combination of results suggests that this subnetwork forms a more specialized module, with increased local segregation reducing its connectivity with the entire network. Our S-Net analysis replicates our L-Net analysis.

These findings are consistent with a recent study by Luo et al. (2024), using a large cross-sectional cohort (n=3355), which demonstrates that unimodal sensorimotor cortices tend to strengthen their connectivity with age, while transmodal association cortices show a decline in connectivity. This pattern highlights fundamental trends in the development of functional connectivity across the cortical hierarchy and supports the idea of a sensorimotor-association axis in the maturation of human cortico-cortical functional connectivity^36^. Our replication of Luo et al.’s findings emphasizes the fundamental role of the SMN in adolescent FBN development^46^.

We also observed a reduction in MBC in L-Net2, which involves parts of the SN, FPN, COTC, and subcortical networks. This suggests that the network tends to reduce functional long-range connectivity with other networks, consistent with a less central role in the overall network structure of this network across mid-adolescent development. In particular, this reduction in integration is not accompanied by an increase in segregation, which is typically seen when specific modules become more distinct, as we observed with SMN. Additionally, L-Net2 is divided into multiple subnetworks in hierarchical analysis, none of which show significant changes in terms of centrality. This suggests that the reduction in MBC for L-Net2 is not due to changes within its individual subnetworks but rather reflects a broader, network-wide shift in its connectivity dynamics.

Finally, we observed a reduction in the MEC of L-Net5, which is mostly encompassed by the VN and the DAN. This reduction suggests a decrease in local connectivity within the module as mid-adolescents age. Since this reduction is not accompanied by changes in MBC or MR, we can infer that the decline in MEC reflects a decomposition of L-Net5 during mid-adolescence. This may indicate that the DAN and VN are becoming more distinct, forming separate modules as mid-adolescent brain development progresses. This finding is further supported by our hierarchical analysis, which shows the emergence of a distinct DAN subnetwork (S-Net4) that becomes separated from the VN over the two-year study.

### 3-2 Grouping based on demographic features

Based on previous studies^4,5,7,8,10,47^ highlighting the significance of CVE and PE in adolescent development, we categorized individuals into groups based on these factors using an unsupervised machine learning approach. CVE has been linked to heightened stress responses, emotional dysregulation, and impaired cognitive function in adolescents^48–53^, as well as to functional and structural neural developmental impacts^6,7,54–56^. PE, on the other hand, has been associated with better access to resources, supportive parenting practices, and enhanced cognitive development in children^57,58^, with corresponding effects on the development of brain structure and function^8–10,48^. Given the complexity of social challenges facing ethnically diverse youth living in low-SES communities, these factors are arguably especially important sources of variability in brain development^7,12,59^.

Our clustering analysis identified three groups, which could be characterized as: high-PE/low-CVE, low-PE/low-CVE, and low-PE/high-CVE. We then examined whether developmental trends differed among these groups, and how psychological measures TT and IQ affect developmental trends within these groups, as we discuss below.

#### 3-2-1 Effects of PE and CVE on development of whole-FBN features

Interestingly, we found distinct trends in developmental changes across the groups. The high-PE/low-CVE group, arguably the most socially advantageously positioned, showed significant changes only in topological FBN integration. The low-PE/low-CVE and low-PE/high-CVE groups exhibited significant changes across all FBN features (topology, dynamics, and complexity). Together these findings suggest greater developmental variation in the high-PE/low-CVE group, compared with more consistent patterns of FBN change among youth whose parents had less education. This convergence toward less flexible and more constrained neural developmental trajectories at the whole-brain level may reflect the influence of environmental stressors, which have been associated with more rigid and less adaptable network configurations^60–64^.

#### 3-2-2 Effects of PE and CVE on development of functional modules

We next examined the development of functional modular changes within the groups. The first observed difference between groups in our modularity analysis pertains to developmental changes within L-Net3, a primary sensory network that includes aspects of both the SMN and the AN. Here, the low-PE/high-CVE group displayed reduced modularity over the two years, indicating that, despite exhibiting more marked changes in whole-FBN features compared to the high-PE/low-CVE group, their SMN-AN network development differed. Supporting this, our hierarchical analysis found that the low-PE/high-CVE group also uniquely showed a nonsignificant developmental trend in S-Net2 (a subnetwork of L-Net3 that includes the AN and lesser aspects of the SMN). At the same time, the low-PE/high-CVE group exhibited a significant developmental trend in S-Net3 (another subnetwork of L-Net3 containing mainly SMN, and not AN). This result was similar to the trend exhibited for the other groups in S-Net3. These results together suggest that high CVE may undermine the formation of a separate auditory sensory network (i.e., AN) as a hierarchical subset of a broader sensory-motor structure. This separation is normative in development from childhood to adulthood^64,65^. Our finding suggests that social stressors may influence the hierarchical organization of somatosensory FBNs.

Second, we noted that the development of modules involving the DMN and SN differed in our sample between the low PE and high PE groups. While the high PE group showed a significant developmental change in the S-Net6, comprised of DMN, the low PE groups showed significant changes in the L-NET2, S-Net9, comprised of aspects of the salience network and subcortical regions, and in the nearby COTC network, anchored in the dorsal anterior insula and dorsal anterior cingulate. This work contributes to literature showing SES effects on functional development in these networks^5,6,10^, and suggests the effects of parental education specifically on FBN development in low-SES mid-adolescents.

### 3-3 Transcendent thinking predicts FBN development

Our results highlight the potential of TT as a resilience factor that influences FBN organization in the face of adversity. In machine learning analyses, TT consistently emerged among the key predictors of complex FBN feature change. In the partial correlation analyses, TT predicted developmental changes in FBN within the low-PE/high-CVE group. Potentially due to higher levels of environmental stresses within this group, longitudinal FBN change in this group was relatively homogenous. And yet, within this group, the two individuals with the highest TT scores were the only participants to demonstrate a different pattern: while others in that group showed positive change in CC and negative change in H, the two participants with highest TT showed negative change in CC and almost no change for H.

Together these results suggest the possibility that TT influences the organization of functional brain connectivity over time, and may be especially helpful under conditions of social stress. This interpretation is consistent with that from our previous work, in which TT predicted young adult psychosocial outcomes via predicting future change in mid-adolescent functional neural connectivity^12^; and in which TT counteracted longitudinal gray matter volume decreases associated with exposure to community violence^7^. Future research should continue to investigate the role of TT in brain development, especially among youth living under disadvantageous social circumstances.

Our findings add to the evidence that TT is a positive factor in mid-adolescent brain development^7,12^, underscoring the lessons from a century of developmental psychological science suggesting the benefits of variants of complex social and identity-related thinking for adolescents^17,66–69^. The potential impact on brain development is notable especially as TT is malleable and can be supported by educational and therapeutic interventions^13^. Future research should continue to examine this factor in the context of brain development, and address the possibility that pedagogies and other structured activities that support dispositions toward TT in mid-adolescents could be beneficial for brain and psychosocial development into young adulthood. These approaches encourage students to grapple with complex ideas and dynamic systems thinking through problem– and project-focused pedagogical approaches and civically framed curriculum^18,59^. There is a growing focus on such approaches to schooling for adolescents, which are associated with a range of psychosocial and academic benefits, especially for youth from underprivileged environments^70–73^.

### 3-4 Conclusion

In conclusion, our study is the first to examine mid-adolescent FBN development longitudinally in a diverse urban sample from a low-SES context. Within our low-SES urban sample, we found consistencies across the group and also developmental effects of social circumstances. Previous studies have shown effects of family income, but financial means are indirect measures of social experience. By separating the effects of parents’ education from those of violence exposure in our low-SES sample, we offer a more nuanced understanding of how two factors highly relevant to adolescents’ daily lives shape FBN development. We also engaged participants in a two-hour open-ended interview that allowed us to gage their spontaneous disposition toward TT, roughly defined as thinking complexly and beyond the “here and now/there and then”^74,75^. We believe that youth with higher scores on our TT measure may be going about their daily lives with more thoughtfulness, reflectivity and social compassion^74^, and that this may over time shape their neural development in a beneficial direction.

We employed a range of advanced methodologies to gain deeper insights into the factors influencing the dynamic development of FBNs in this population. We found that participants whose parents had lower levels of education underwent more pronounced changes in whole-FBN characteristics. In our modularity analysis, while CVE was associated with differences in somatosensory and auditory network development, PE was associated with influences on DMN and salience network development. TT emerged as a key predictor of FBN development across the sample, and particularly in the context of high CVE. Together our findings contribute new and potentially actionable insights into the sources of variability in FBN development among mid-adolescents living in low-SES environments.

### 3-5 limitations

Given the exploratory nature of this study—the first of its kind—a comprehensive analysis with a modestly-sized sample was necessary. The generalizability of our findings may be revealed with a larger sample-size in future; we note that TT predicted FBN development across all groups before correction for multiple comparisons. Given our promising results, future studies are needed in particular to investigate how resilience factors like TT may support neural plasticity and buffer the effects of environmental stressors in a range of social contexts.

## 4– Methods

The data used in this study were drawn from a broader research project in which participants also took part in additional psychosocial tasks, psychophysiological assessments, and neuroimaging procedures not directly related to the current analyses (e.g., interviews about school experiences, heart rate variability studies, diffusion tensor imaging; see https://osf.io/gqs34 for details). All procedures were reviewed and approved by the University of Southern California Institutional Review Board (UP-12-00206) and conducted in accordance with its ethical guidelines. Written informed consent or assent was obtained from all participants and their parents or legal guardians, as appropriate. Participants were compensated for their time.

### 4-1 Participants

Sixty-five healthy, right-handed healthy mid-adolescents (age 14-18 years at recruitment, M_age_=15.8 years, SD=1.1 years; 36 female/29 male) were recruited to participate in the initial session from public high schools in low-SES neighborhoods with moderate to high levels of crime in Los Angeles. Participants were from stable families, passing all their classes at school, with no diagnosed psychiatric issues or developmental or learning disabilities, not using drugs or alcohol, and not under disciplinary action in or outside of school. Enrollment criteria included no history of: neurological or psychiatric disorders; physical or emotional abuse or neglect; use of psychotropic medication, recreational drugs or alcohol; a medical condition that would preclude scanning. Receiving free or reduced-price lunch, an indicator of low SES due to a low family income-to-needs ratio, was reported by 51 participants. To ensure a culturally/ethnically diverse sample, at least one of each participant’s parents was born and raised to adulthood outside of the U.S. (participants’ parents originated from 13 countries). Additional criteria required participants to be enrolled in school full-time, passing all classes, fluent in English, and not subject to any disciplinary actions. None had been directly involved in violent crimes, either as victims or perpetrators.

Roughly two years later (M_time between visits_=2.09 years, SD=0.21 years), 62 of the 65 participants returned for the second laboratory visit. (Attrition resulted from participants moving out of the region.) 6 participants were not able to undergo MRI scanning at the second visit due to: dental braces (4 participants); metal pins in the leg (1 participant); or a technical problem with the scanner (1 participant).

### 4-2 Procedures and Acquisition of Psychosocial and Demographic Measures

#### 4-2-1 Psychosocial Interview Capturing Transcendent Thinking (TT)

As described in Gotlieb et al., 2022 and 2024^15,74^, at the initial session, participants reacted to 40 true, compelling stories about living, non-famous adolescents from around the world in a range of circumstances, during a two-hour private video-taped interview (the protocol was adapted from Immordino-Yang et al., 2009^77^). The story corpus had been previously piloted to ensure it was engaging and capable of eliciting a range of both positive and negative emotional responses. The experimenter recounted each story using a memorized script, then played a one-minute, documentary-style video featuring real-life footage of the actual protagonist involved in the story (not an actor). Videos were presented using PowerPoint (Microsoft Office) on a 17-inch Lenovo laptop. After each video, the experimenter asked the participant, ‘How does this story make you feel?’ and then looked down to take handwritten notes, attempting to capture the participant’s response as accurately as possible. Participants were told the notes were a precaution in case of video recording failure. In fact, they also served to standardize the experimenter’s behavior to facilitate free responses from the participants. Participants were encouraged to as candid as possible.

As reported in Gotlieb et al. 2024^74^, videotaped interviews were transcribed and verified. Participants’ responses were blind-coded and reliability coded to identify constuals reflecting TT. Such construals were defined as utterances reflecting:

(i) systems-level evaluations or moral judgements, or curiosities about how and why systems work as they do, e.g.,

> “I also find it unfair that the people get undocumented. It’s kind of weird how it’s like a label how like just ‘cause you are from some other place, um, you can’t do certain things in another place. It’s like a question. It’s like something I’ve always wondered…”;
(ii) discussions of broad implications, morals and moral emotions, perspectives, personal lessons or values derived from the story, e.g.,

> “I think back to the idea that because children are the future […] we have to be able to inspire people who are growing and have the potential to improve the societies”;
>
> “it makes me happy for humanity”;

or (iii) analyses of the protagonist’s qualities of character, mind, or perspective, e.g.,

> “[she is] thinking, ‘oh, you’re not alone. You have others who are dependent on you’.”

Importantly, it was not relevant whether the participant endorsed a value or lesson or agreed with the protagonist, e.g.,

> “I wouldn’t react that way. I’d just be really mad at the kid instead of, you know, selfless like that and trying to help him. Like I wouldn’t be able to put myself in someone’s shoes like that like he did.”

Construals that were not identified as reflecting TT primarily centered on the protagonist’s immediate circumstances, e.g., “I’m glad it all worked out,” or involved judgments about the protagonist’s choices or behaviors, such as “I feel like they should have planned it more.” Others reflected the participant’s empathic emotional responses, e.g., “I feel really sad for her, and like, second-hand embarrassment.” In contrast to TT, these responses are characterized by concrete, context-specific, and reactive interpretations.

Participants received a score of 1 for each construal indicating TT. Scores across all trials were summed to produce a total score for each participant.

#### 4-2-2 Parental education (PE)

As described in Yang et al., 2025^7^, at session 1, participants reported the highest level of education achieved by their parents, a factor associated with SES^8^. Following work by Noble et al. (2015), PE level was recoded as years of formal education, as follows: did not complete high school = 8 years; received a high school diploma or a General Education Development (GED) degree=12 years; some college/received an Associate’s degree or postsecondary vocational certificate = 14 years; received a Bachelor’s degree = 16 years; received a Master’s or a Doctoral degree = 18 years. PE data were unavailable for four Latino participants. For these participants, parental years of education were estimated based on the average years of schooling (10 years) reported by Latino participants from the same neighborhood.

#### 4-2-3 Community Violence Exposure (CVE)

As described in Yang et al., 2025^7^, at sessions 1 and 2, participants reported their CVE using a modified 13-item version of the Survey of Children’s Exposure to Community Violence – self-report version^78^. The questionnaire includes a list of thirteen events that involve either threatened or actual victimization (e.g., Item 9. Has anyone ever threatened to beat you up?) or witnessing and/or hearing about violence (e.g., Item 13. Have you ever witnessed or heard about someone being shot?). For each item, participants chose from four options, including “Never,” “Once,” “Twice” or “More than twice.” At session 1, participants were instructed to report any incidents of violence exposure they remembered experiencing (to measure childhood and early adolescent CVE, prior to participation in the study). At session 2, they were instructed to report new incidents over the two-year period since the first session (to measure mid-adolescent CVE).

Following the questionnaire, each participant took part in a private interview to verify their responses and to identify whether the reported exposure to violence happened within the community (neighborhood or school), through media sources (e.g., television), or at home. Incidents observed through media were excluded from the violence exposure score.

Participants would have been disqualified if incidents occurred in the home or involved family members; however, no such cases were reported.

For each item, participants received a score from 0 (“Never”) to 3 (“More than twice”), giving a potential final score of between 0 and 39. Total scores were separately tallied for childhood/early adolescent CVE (reported in session 1) and mid-adolescent CVE (reported in session 2).

#### 4-2-4 IQ testing

As described in Gotlieb et al. 2024^74^, during the second session, a trained experimenter individually conducted the vocabulary and matrix reasoning subtests from the second edition of Wechsler Abbreviated Scale of Intelligence^79^, in a private room. Scores from these subtests were age-normed, combined, and totaled to yield an overall IQ score for each participant. Due to time limitations, one participant completed only the verbal subtest, and their overall IQ was imputed based on this score.

The psychological and demographic measures for all participants are available at: https://osf.io/9jxaz/.

### 4-3 MRI acquisition

As described in Gotlieb et al.,2022^15^, at sessions 1 and 2, following acquisition of psychosocial measures, participants underwent a seven-minute resting-state BOLD fMRI scan. We instructed participants to think about whatever they wished, remain as still as possible, and stay awake. We continuously displayed an image of a nature scene without any people or animals. We scheduled the protocol so that scans would take place around midday.

BOLD fMRI data during the session 1 were collected using a 3 Tesla Siemens Trio scanner equipped with a 12-channel matrix head coil. A T2∗-weighted echo-planar imaging (EPI) sequence was used to acquire the functional scans (TR = 2 s, TE = 25 ms, flip angle = 90°, acquisition matrix = 64 × 64, FOV = 192 mm), with a voxel size of 3 × 3 × 3 mm³. Forty-one interleaved transverse slices were obtained to provide whole-brain coverage. In total, 210 functional volumes were collected. High-resolution anatomical images were acquired using a magnetization-prepared rapid gradient echo (MPRAGE) sequence (TI = 800 ms, TR = 2530 ms, TE = 3.09 ms, flip angle = 10°, isotropic voxels of 1 mm³, acquisition dimensions of 256 × 256 × 176).

At session 2, due to a system upgrade at the scanning facility, imaging was conducted using a 3 Tesla Siemens Prisma scanner equipped with 20-channel matrix head coils. The parameters for BOLD fMRI acquisition remained unchanged from the session 1. For anatomical imaging, we used an MPRAGE sequence (TI = 800 ms, TR = 2530 ms, TE = 3.13 ms, flip angle = 10°, isotropic voxel resolution of 1 mm³, acquisition dimensions of 256 × 256 × 256).

Analyses were conducted to ensure that our effects are not due to the scanner upgrade; see *Preparatory analyses* section below.

### 4-4 MRI data Preprocessing

MRI data preprocessing was performed using fMRIPrep 20.0.7 (RRID:SCR_016216)^80^, which is based on Nipype 1.4.2^80,81^ (RRID:SCR_002502).

#### 4-4-1 Anatomical data preprocessing

T1-weighted (T1w) images from both sessions 1 and 2 were corrected for intensity non-uniformity (INU) with N4BiasFieldCorrection^82^, distributed with ANTs 2.2.0^83^ (RRID:SCR_004757). The T1-weighted reference image then underwent skull stripping using the Nipype implementation of the antsBrainExtraction.sh script (from ANTs), with the OASIS30ANTs template serving as the target. Brain tissue segmentation of cerebrospinal fluid (CSF), white-matter (WM) and gray-matter (GM) was performed on the brain-extracted T1w using fast (FSL 5.0.9, RRID:SCR_002823^84^). A T1w-reference map was computed after registration of 2 T1w images (after INU-correction) using mri_robust_template (FreeSurfer 6.0.1)^85^. Volume-based spatial normalization to a standard space (MNI152NLin6Asym) was performed through nonlinear registration with antsRegistration (ANTs 2.2.0), using brain-extracted versions of both T1w reference and the T1w template. The following template was utilized for spatial normalization: FSL’s MNI ICBM 152 non-linear 6th Generation Asymmetric Average Brain Stereotaxic Registration Model (RRID:SCR_002823; TemplateFlow ID: MNI152NLin6Asym)^86^.

#### 4-4-2 Functional data preprocessing

The following preprocessing was performed for each resting-state scan. Using a custom fMRIPrep methodology, a reference volume and its skull-stripped version was first generated. Susceptibility distortion correction (SDC) was omitted. The BOLD reference was then co-registered to the T1w reference using flirt (FSL 5.0.9)^87^, with the boundary-based registration^88^ cost-function. Co-registration with nine degrees of freedom was applied to correct for any remaining distortions in the BOLD reference image. Using mcflirt (FSL 5.0.9)^87^, the head motion parameters in relation to the BOLD reference image (transformation matrices and six motion components of translations and rotations) were estimated before applying any spatiotemporal filtering. For slice-timing correction of the BOLD runs, 3dTshift from AFNI version 20160207 (RRID:SCR_005927) ^89^ was used. After that, we resampled the BOLD time series, including any slice-timing corrections, back into each participant’s native space by applying the motion correction transforms. We refer to these resampled series as preprocessed BOLD in native space, or simply preprocessed BOLD. The BOLD time-series were resampled into a standard space (MNI152NLin6Asym), generating spatially-normalized, preprocessed BOLD runs. Several confounding time-series were calculated based on the preprocessed BOLD: framewise displacement (FD), DVARS, and three region-wise global signals. For each functional run, FD and DVARS were computed using Nipype tools, based on the criteria outlined by Power et al. (2014)^90^. Global signals from CSF, WM, and the entire brain volume were extracted using their corresponding masks.

#### 4-4-3 Removal of motion artifacts and other confounds

Motion artifacts were removed from the preprocessed BOLD time-series in MNI space using an independent component analysis based strategy for automatic removal of motion artifacts (ICA-AROMA)^91^, as implemented in fMRIPrep 20.0.7. This procedure was applied following the exclusion of non-steady state volumes and after spatial smoothing with a 6 mm full-width at half-maximum (FWHM) isotropic Gaussian kernel. Corresponding “non-aggressively” denoised runs were generated post-smoothing. The spatially-normalized “non-aggressively” denoised resting state BOLD time-series were further processed using XCPEngine^92^ to apply additional steps in the ICA-AROMA procedure detailed in Pruim et al., 2015^91,93^. Specifically, BOLD time series were temporally filtered using a first-order Butterworth filter with a passband between 0.01 and 0.08 Hz. A confound regression step additionally regressed out mean white matter and cerebral spinal fluid signals. In order to prevent frequency-dependent mismatch during confound regression^94^, all regressors were band-pass filtered to retain the same frequency range as the data.

### 4-5 Functional connectivity and brain network construction

Using XCPEngine, we applied a commonly used whole brain parcellation consisting of 264 spherical nodes containing cortical and subcortical brain regions^46^. For each region, the mean time series across all voxels was calculated from the denoised residual data. Data from 12 brain regions were excluded from analysis because they were missing for at least one participant. As a result, time series data from 252 brain regions were used for FBN construction. The list of excluded regions is presented in supplementary table 1.

To model functional brain networks, activity of parcellated brain regions were considered as “nodes” or vertices; coherence between nodes were defined as “edges” or functional connections^95^. We calculated coherence between BOLD signals of all brain regions (separately for individuals and sessions). To do this, magnitude squared coherence between signals *x* and *y* were computed using MATLAB R2024a’s *mscohere* function (*C_xy_* = *mscohere*(*x,y*)). This function estimates coherence via Welch’s method, which involves averaging modified periodograms. Coherence quantifies the frequency-domain associations between *x* and *y*, ranging from 0 to 1, where higher values indicate stronger alignment at a given frequency. The magnitude squared coherence (*C_xy_*) is defined as^95^:

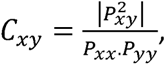

where *P_xx_* and *P_yy_* are the Power Spectral Density (PSD) estimates of *x* and *y*, respectively, and *P_xy_* represents the Cross Power Spectral Density (CPSD) estimate of *x* and *y*. By default, *mscohere* partitions *x* and *y* into eight 50% overlapping segments, applies a Hamming window to each segment, and averages the resulting periodograms to enhance estimation accuracy. Then, we average across all frequencies and use the average values as the weighted undirected connections to construct adjacency matrices^96^. In an adjacency matrix, the cell at the intersection of the *i-th* row and *j-th* column represents the edge between the *i-th* and *j-th* nodes of the network; the weight of the edge is stored in this cell. One matrix for each participant for each session was produced. Weighted undirected adjacency matrices were then used for further network analyses. All adjacency matrices are available at https://osf.io/9jxaz/.

### 4-6 Brain network analysis

#### 4-6-1 Whole brain network features

Adjacency matrices were analyzed to reveal FBN properties. First, we calculated the average connectivity between all nodes and considered this measure as an index of connection density (CD). This measure reflects the overall level of connectivity in the network, regardless of the topology of the paths or the synchronization and dynamics of the nodes. We then proceeded to analyze complex network measures from graph theoretical analysis. To assess the topological features present in FBNs, we used two topological measures: CC and CPL, which provide information about the architecture of paths between nodes^38,96,97^, as follows:

The CC reflects the local processing and functional segregation in FBNs. Networks with highly interconnected triple nodes (triangle), considered as clusters, exhibit high values of CC. Given the brain’s modular mechanisms for neural signal processing, functional brain networks are hypothesized to be segregated networks^38,96,97^; CC can be used to quantify this segregation. For a weighted undirected network, CC is defined by^98^:

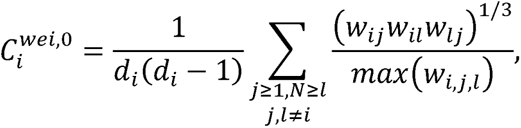

where *d_i_* is the degree of node *i*, *w_ij_* is the connectivity weight between nodes *i* and *j*, and max(*w_i,j,l_*) equals to the maximum weight among the three nodes in the triangle. To calculate this measure, we used *clustering_coef_wu* function from the Brain Connectivity Toolbox^96^.

The CPL quantifies the information integration that arises from distributed nodes within the FBN. This measure is related to the average of all shortest path lengths between pairs of nodes in the network. Networks with lower shortest path lengths between node pairs (lower CPL) exhibit a higher capacity for fast information integration. To calculate CPL for a weighted undirected network, we first inverted all the connectivity weights to convert them from connection values between nodes to path length values. Then, we applied the *distance_wei_floyd* algorithm to compute CPL over FBNs. This algorithm was implemented in the Brain Connectivity Toolbox^96^.

To evaluate the dynamics of coupling between all pairs of nodes, we employed spectral graph theoretical analysis^38^. In spectral graph theoretical analysis, energy (H) is associated with the stability of synchronization in the network. To calculate H, we calculate the eigenvalues of each adjacency matrix (weighted and symmetric) and H is equal to^38^:

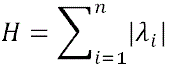

where λ*_i_* is eigenvalue of node *i*, and *n* is number of nodes. We computed H using a self-developed MATLAB 2024a function available at https://github.com/AHGhaderi/Amir-Hossein-Ghaderi/commit/614da0f285b47a64ac501df0606cd6b4cf665460.

Finally, from information theory, we employed Shannon entropy (S), which reflects the complexity of the functional connectivity pattern between nodes^38^. S is defined by:

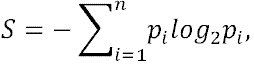

where *p_i_* is the probability of occurrence of *i^th^* connection in the adjacency matrix and *n* is the number of connections. S is inversely associated with the probability of state occurrence in a system. For example, minimum S is achieved when all connections (or states) among nodes are identical, meaning that the probability of that state is 1. Higher values of S occur when rare connectivity values exist among the connections or when the distribution of connectivity values is more random. To calculate S for FBNs, we used a self-developed MATLAB 2024a code, which is available at https://github.com/AHGhaderi/Amir-Hossein-Ghaderi/commit/db17d96d35b73e8eee8fa526ae307a457ad62272.

We also applied null models to normalize our graph measures. The null models were generated 1000 times by randomly permuting the edge weights while preserving the degrees of nodes and matrix symmetry^42^. Specifically, we divided the CC, CPL, and H by their corresponding average values computed from the null models. Since CD and S are not dependent on the spatial arrangement of elements in the adjacency matrix, their values remained unchanged between the original and null model matrices. To generate null models, we used self-developed MATLAB 2024a codes available at: https://github.com/AHGhaderi/Null-model.

#### 4-6-2 Modularity analysis and subnetworks

For modularity analysis, we employed a combination of unsupervised and supervised modularity detection approaches. An unsupervised algorithm can identify clusters of nodes that are strongly connected to each other^43,44^. However, this algorithm includes a *modularity parameter* that can regulate the strength of connectivity, thereby increasing or decreasing the number of clusters (see the *modularity_und* function at Brain Connectivity Toolbox^96^). Typically, allowing clusters to form at lower connectivity strengths results in larger clusters, leading to a network division into fewer segments. Conversely, when stricter thresholds are applied, requiring clusters to exhibit higher connectivity strengths, the clusters tend to become smaller, and the network is divided into a larger number of small subnetworks. To determine the appropriate *modularity parameter*, we employed a supervised modularity approach. This analysis consists of the following steps^45^:

1. Choose a minimum value for the *modularity parameter* in the unsupervised algorithm (*modularity_und* function), ensuring it reveals at least two subnetworks.
2. Run the unsupervised modularity analysis.
3. Select the group of nodes (i.e., brain regions) identified as a subnetwork by the unsupervised algorithm.
4. Compute the average connectivity among the selected nodes.
5. Randomly shuffle connections across the entire network.
6. Compute the average connectivity among the selected nodes after randomization.
7. Calculate the *modularity ratio (MR)* by dividing the connectivity average from Step 4 (original subnetwork) by the connectivity average from Step 6 (randomized network).
8. Repeat Steps 3 to 7 for all subnetworks and average the *MR* values obtained in Step 7.
9. Increase the *modularity parameter* value and repeat Steps 2 to 8 until the network is divided into isolated nodes instead of subnetworks.
10. Identify the maximum averaged *MR* from Step 8 and determine the corresponding *modularity parameter* for this ratio.

For consistent analysis across all networks, we used an averaged adjacency matrix derived from all individual adjacency matrices including both sessions. Once the optimal modularity parameter was identified, it was used for all subsequent analyses. The functions for this analysis were implemented in MATLAB 2024a; self-developed open access code is available at https://github.com/AHGhaderi/Amir-Hossein-Ghaderi/commit/df636b2105e578a8969ffa88856e54f3267a40a7.

We also introduced and utilized two novel parameters in this study: modular eigenvector centrality (MEC) and modular betweenness centrality (MBC). To calculate *MEC*, we first computed the eigenvector centrality for each node within a given module and then averaged these values. The eigenvector centrality for each node was derived by calculating the eigenvalues and eigenvectors of the adjacency matrices. Specifically, for each given adjacency matrix, the eigenvector corresponding to the largest eigenvalue was considered as the eigenvector centrality of the respective nodes. The eigenvector centrality calculation for each node was performed in MATLAB 2024 using *eigenvector_centrality_und* function in Brain Connectivity Toolbox^96^. To calculate *MBC*, we followed a similar approach by first computing the betweenness centrality for each node within a given module and then averaged these values. The betweenness centrality of each node was calculated by counting the shortest paths passing through that node, implemented using the *betweenness_wei* function from the Brain Connectivity Toolbox^96^.

To ensure that the observed values of MEC and MBC were not driven by overall connectivity average in the network, again we applied null model corrections^42^. Specifically, we generated shuffled networks preserving the original degrees on nodes and for each shuffled network, we computed MEC and MBC using the same procedure as for the original network. We then normalized the observed values by subtracting them from the values of shuffled networks.

### 4-7 Data analysis for hypothesis testing

#### 4-7-1 Preparatory analyses

Prior to conducting the main analyses, we carried out a preliminary investigation to examine potential sex-related differences in network measures. Participants were categorized by sex, and FBN features were compared between females and males using t-tests. No statistically significant differences were found (Supplementary Figure 1).

To ensure that the MRI system upgrade that took place between sessions 1 and 2 did not influence the FBN metrics, we identified maximum age from the first session and minimum age from the second session and compared FBN measures from the 42 participants who fell within this range (i.e., we compared participants of the same age when they were scanned with either the older or the upgraded scanner). In the between-subject statistical analysis, 6 participants for whom both sessions fell within the identified age range were excluded from the second scan group. No significant differences were observed between sessions 1 and 2 across any FBN measures (see also Supplementary Figure 2).

#### 4-7-2 Group classification with k-mean analysis

To cluster participants into different groups, we used k-means analysis based on parental education (PE) and community violence exposure (CVE). K-means clustering is an unsupervised machine learning algorithm that partitions data into k distinct groups by minimizing within-cluster variance, iteratively assigning each data point to the nearest centroid. To determine the optimal number of clusters, we employed the Calinski-Harabasz index (also known as the variance ratio criterion)^99^, a metric that evaluates cluster quality by calculating the ratio of between-cluster dispersion and within-cluster dispersion; higher values of this index indicate better-defined clusters. For robustness, we tested multiple values of k and selected the solution with the highest Calinski-Harabasz score. K-means clustering was performed using MATLAB’s default implementation (*kmeans* function) with the following parameters: Distance metric: Squared Euclidean; initialization: k-means++ for improved centroid seeding; maximum iterations: 100; and replicates: 10 (to avoid local minima). The algorithm converged when centroid movements fell below 1^e-4^.

#### 4-7-3 Permutation t-tests

We performed within-subject comparisons of network features between session 1 and session 2 using nonparametric paired permutation t-tests. The permutation t-test is a robust, distribution-free method for assessing statistical significance by randomly shuffling condition labels within subjects to generate a null distribution of the test statistic^100^. This approach maintains the dependence structure of repeated-measures designs and avoids assumptions of normality, making it particularly well-suited for neuroimaging and network-based analyses. For each network feature, we computed the observed t-statistic across participants, then randomly permuted the condition labels (session 1 vs. session 2) within each subject 5,000 times. For each permutation, a new t-statistic was computed, forming an empirical distribution that approximates the null distribution of t-values. With 5,000 permutations, this null distribution tends to approach a normal shape due to the central limit theorem, enabling reliable estimation of p-values. The p-value was determined by the proportion of permuted t-statistics that were greater than or equal to the observed value in absolute magnitude. To address multiple comparisons, we conducted separate false discovery rate (FDR) corrections (Benjamini-Hochberg procedure)^101^ for global network features (i.e., CD, CC, CPL, H, and S) as well as modularity-specific characteristics (i.e., MR, MBC, and MEC).

We implemented paired permutation t-tests using custom MATLAB functions available at https://github.com/AHGhaderi/Amir-Hossein-Ghaderi/commit/267f8633d1fb728cfe54f2984f9a84f258475070. FDR analysis was performed using *fdr_bh* function in MATLAB 2024a available at https://www.mathworks.com/matlabcentral/fileexchange/27418-fdr_bh.

#### 4-7-4 Longitudinal analyses

##### 4-7-4-1 Linear mixed model

To evaluate longitudinal changes in network features, we employed linear mixed-effects models with age as a fixed effect in each analysis. These models were designed to account for within-subject dependencies across sessions while assessing developmental trajectories. Importantly, we included random intercepts for participants to accommodate individual variability in baseline measures. This modeling choice was particularly important because not all participants were present in every session; some contributed data from only the first session. The inclusion of random intercepts allows the model to appropriately handle such unbalanced data and ensures that the variability attributable to individual differences does not confound the estimated fixed effects. To independently assess the developmental trajectory of each network property, separate linear mixed-effects models were estimated for each network feature. The general structure of the model was as follows:

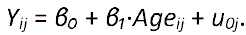

Where: *Y_ij_* is the network feature value for subject *j* at time point *i*, *β₀* is the fixed intercept, *β*₁ is the fixed effect of age, *u_0j_* is the random intercept for subject *j*. To account for multiple comparisons across network features, we applied FDR adjustment. All reported *q*-values represent FDR-corrected *p*-values, thereby controlling Type I error rates.

##### 4-7-4-2 Machine learning prediction, F-test predictors ranking and SVM

To further validate our previous findings and assess the effects of various demographic and psychological measures on predicting developmental patterns for each individual, we applied an *F*-test feature selection analysis followed by supervised support vector machine (SVM) regression. These analyses were implemented using MATLAB’s Machine Learning app (R2024a).

The goal was to predict whole-brain network measures in the second session (the target variable) based on FBN measures from the first session and psychological/demographic variables (TT, IQ, PE, and CVE), as well as age at both sessions.

*F*-test feature ranking was applied to evaluate the statistical significance of each predictor’s linear relationship with the target variable. MATLAB’s default *F*-test method calculates the *F*-statistic as the ratio of the between-group variance to within-group (residual) variance for each feature independently; features were ranked accordingly.

Then, we used the Fine Gaussian SVM regression model, using an ε-SVM regression model with a radial basis function (RBF) kernel. The following default parameters were applied: kernel function: Gaussian (RBF); kernel scale: automatically optimized using heuristics based on the distribution of predictor values (MATLAB default: ‘auto’, using a variant of the median heuristic); box constraint (C): 1; epsilon (ε): 0.1; standardization: enabled (predictors were z-scored prior to training); solver: iterative SMO (Sequential Minimal Optimization). Model performance was evaluated using 5-fold cross-validation to ensure robustness and generalization. As an accuracy metric, we calculated the ratio of the standard deviation of the true (observed) target values to the model’s mean squared error (MSE). A higher ratio indicates greater predictive accuracy, with the model accounting for a larger portion of the variance in the outcome variable.

##### 4-7-4-3 Partial correlation analysis

Additionally, we performed a partial correlation analysis using MATLAB’s built-in *partialcorr* function to directly assess the relationship between psychological variables and changes in FBN measures. In this analysis, the effect of age at both sessions was controlled. The partial correlation coefficient between each FBN measure and each psychological variable (TT, IQ) was calculated. FDR correction was applied.

## Acknowledgments

We are grateful to L.G. Cardona, D. Cremat, E. Jahner, L. Kim, C. Kundrak, H. Rajana, R. Riveros, and C. Simone for their support with data collection and processing, and to P. Baniqued for analytical guidance. We also thank Artesia High School (Lakewood, CA), Rowland Unified School District (Rowland Heights, CA), and other participating public high schools for their assistance with participant recruitment. This research was supported by National Science Foundation grants (CAREER 11519520; BCS 1522986), a Raikes Foundation grant (61405837-118286), and philanthropic contributions from the ECMC Foundation and Stuart Foundation to M.H.I.Y.

