## Supplementary material for "Intersecting effects of social circumstances and transcendent thinking on mid-adolescents’ longitudinal functional connectome development"

Supplementary Table 1. Excluded regions.

|  | 711-2L space | | | MNI space | | | Suggested network |
| --- | --- | --- | --- | --- | --- | --- | --- |
| **ROI** | x | y | z | x | y | z |  |
| **4** | -53 | -45 | -24 | -56 | -45 | -24 | Uncertain |
| **10** | 50 | -36 | -24 | 52 | -34 | -27 | Uncertain |
| **21** | 26 | -21 | 69 | 29 | -17 | 71 | Sensory/somatomotor Hand |
| **27** | -38 | -30 | 66 | -38 | -27 | 69 | Sensory/somatomotor Hand |
| **37** | -38 | -18 | 66 | -38 | -15 | 69 | Sensory/somatomotor Hand |
| **38** | -17 | -48 | 69 | -16 | -46 | 73 | Sensory/somatomotor Hand |
| **116** | 62 | -15 | -15 | 65 | -12 | -19 | Default mode |
| **127** | 28 | -76 | -31 | 28 | -77 | -32 | Default mode |
| **182** | -20 | 36 | -15 | -21 | 41 | -20 | Uncertain |
| **184** | 17 | -79 | -34 | 17 | -80 | -34 | Uncertain |
| **249** | 47 | -6 | -33 | 49 | -3 | -38 | Uncertain |
| **250** | -47 | -9 | -36 | -50 | -7 | -39 | Uncertain |

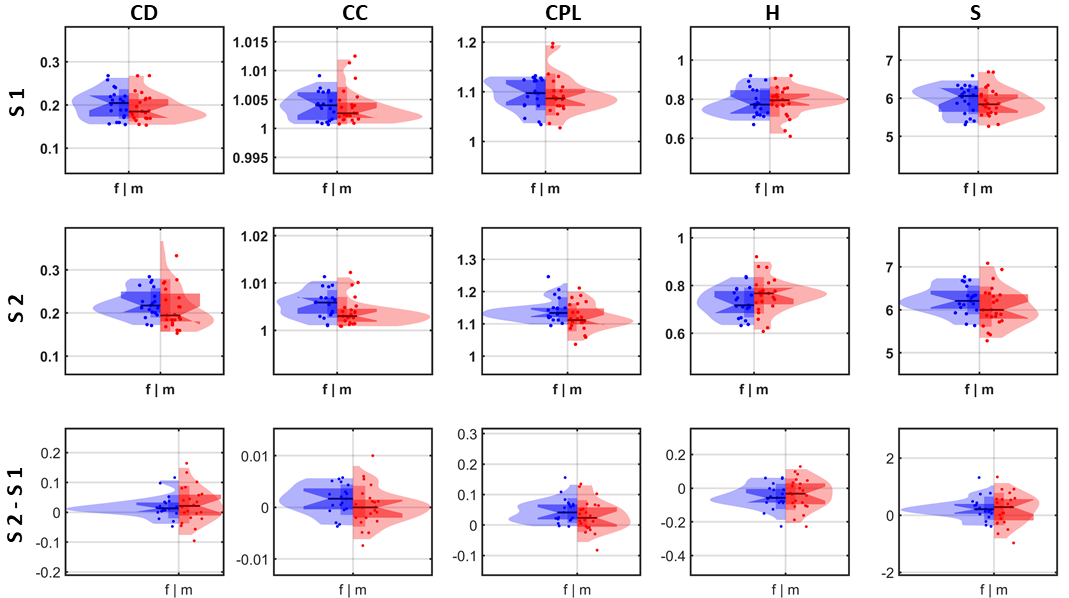

Supplementary Figure 1. Comparison of FBN features between female (f) and male (m) participants in session 1 (S1), session 2 (S2) and for the difference between sessions (S2-S1). No significant difference in FBN features between the two groups was observed.

**S1:** CD: p-value=0.477, CC: p-value=0.971, CPL: p-value=0.991, H: p-value=0.878, S: p-value=0.590.

**S2:** CD: p-value=0.708, CC: p-value=0.093, CPL: p-value=0.064, H: p-value=0.180, S: p-value=0.222.

**S2-S1:** CD: p-value=0.683, CC: p-value=0.261, CPL: p-value=0.239, H: p-value=0.625, S: p-value=0.857.

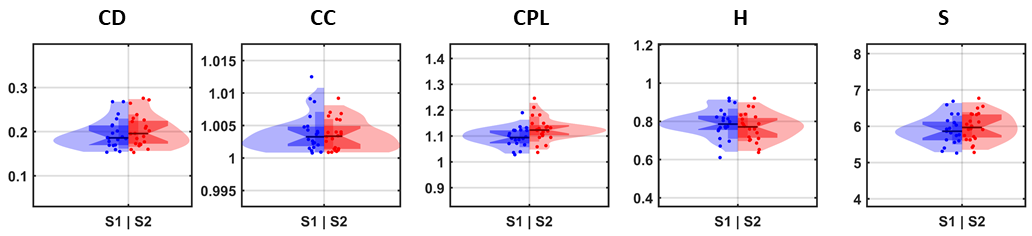

Supplementary Figure 2. Comparison of FBN features between different participants within the same age range scanned before versus after the scanner upgrade, i.e., in session 1 (n = 20, age range = 16.25 to 18.67, mean = 16.98) versus in session 2 (n = 22, age range = 16.11 to 17.90, mean = 17.12). No significant differences were observed between the two groups in age (p = 0.471) or FBN features (CD: p-value=0.543, CC: p-value=0.615, CPL: p-value=0.144, H: p-value=0.479, S: p-value=0.448), suggesting no significant effect of the scanner upgrade on FBN features.
